# Microrheology reveals microscale viscosity gradients in planktonic systems

**DOI:** 10.1101/2020.04.09.033464

**Authors:** Òscar Guadayol, Tania Mendonca, Mariona Segura-Noguera, Amanda J. Wright, Manlio Tassieri, Stuart Humphries

## Abstract

Microbial activity in planktonic systems creates a dynamic and heterogeneous microscale seascape that harbours a diverse community of microorganisms and ecological interactions of global significance. In recent decades a great effort has been put into understanding this complex system, particularly focusing on the role of chemical patchiness, while overlooking a physical parameter that governs microbial life and is affected by biological activity: viscosity. Here we use microrheological techniques to measure viscosity at length scales relevant to microorganisms. Our results reveal the viscous nature and the spatial extent of the phycosphere, the microenvironment surrounding phytoplankton cells, and show heterogeneity in viscosity at the microscale. Such heterogeneity affects the distribution of chemicals and microorganisms, with pervasive and profound implications for the functioning of the planktonic ecosystem.

**One Sentence Summary:** Microrheology measurements unveil the existence of layers of increased viscosity surrounding phytoplankton cells and within aggregates.

## Main Text

Planktonic microorganisms inhabit a microenvironment far more complex and dynamic than suspected even two decades ago. Microbial activity, advection and diffusive transport together create a patchy chemical landscape that within a few microlitres sustains a diverse community of microalgae, protists, bacteria and viruses, along with an intricate network of individual interactions and ecological processes of global significance (*1-3*). However, viscosity, a dominant feature of microbial life, has been overlooked in this emerging paradigm of microscale heterogeneity. Viscous forces govern directed motion and diffusion of microorganisms, particles and molecules in the very low Reynolds numbers regime experienced by microbes. This implies that small microorganisms may not directly experience turbulent flows, that biomechanical plans working for large organisms become inefficient for microbes and that mass transfer is ultimately governed by molecular diffusion, which is inversely proportional to viscosity. Several studies [compiled in (*4*)] have reported increases, often higher than one order of magnitude, of the bulk-phase viscosity of the water due to the presence in solution of extracellular polymeric substances (EPS) exuded by phytoplankton and bacteria. Moreover, it is known that intracellular viscosities can be up to two orders of magnitude higher than water (*5*). However, very little is known about the role viscosity plays in shaping the heterogeneous microscale landscape.

Here, we hypothesize the existence of microscale viscosity gradients created by the same biological processes that generate chemical gradients around planktonic microorganisms (*2, 6*). We argue that the existence of microscale patchiness in viscosity has fundamental implications for the distribution of microorganisms and their resources, and therefore for the functioning of microbial planktonic ecosystems driving aquatic productivity globally.

To test our hypothesis, we characterized the variability in viscosity near microorganisms, non-destructively and at micron length scales, by employing two methodologies from the fast-evolving field of microrheology: microrheology with optical tweezers (MOT) and multiple particle tracking microrheology (MPTM). We investigated three distinct scenarios where we expected viscosity to be heterogeneous: (i) the phycosphere around healthy EPS-exuding phytoplankton cells (*6*); (ii) lysis events involving the release of cell content and (iii) algal cell aggregates.

To validate our hypothesis, we first took advantage of the high spatiotemporal resolution of MOT to measure viscosity near glass shards, as an inert control that mimics siliceous frustules, and near cells of *Chaetoceros affinis*, a diatom known to produce EPS and mucilages (*7, 8*) that lead to the formation of transparent exopolymer particles (TEP). Our results reveal that the relative viscosity (defined as the ratio of local dynamic viscosity to the dynamic viscosity of artificial seawater) is often much higher and more variable near the cell wall of diatoms than near the glass shards. In the case of the diatoms, the relative viscosity is frequently much higher than predictions by Faxén’s law, which accounts for hydrodynamic interactions of a sphere near a solid wall (Fig. 1A, see also Materials and Methods). Moreover, for viscosity measurements taken near the diatoms, we also detected anisotropy in the observations recorded along the axes perpendicular and parallel to the cell wall (Fig. 1B), suggesting that EPS might be discreetly patterned in space. These observations are consistent with studies showing the presence of mucilaginous or gel-like structures around fixed bacteria and phytoplankton exuding EPS (*9-12*).

**Fig. 1:**
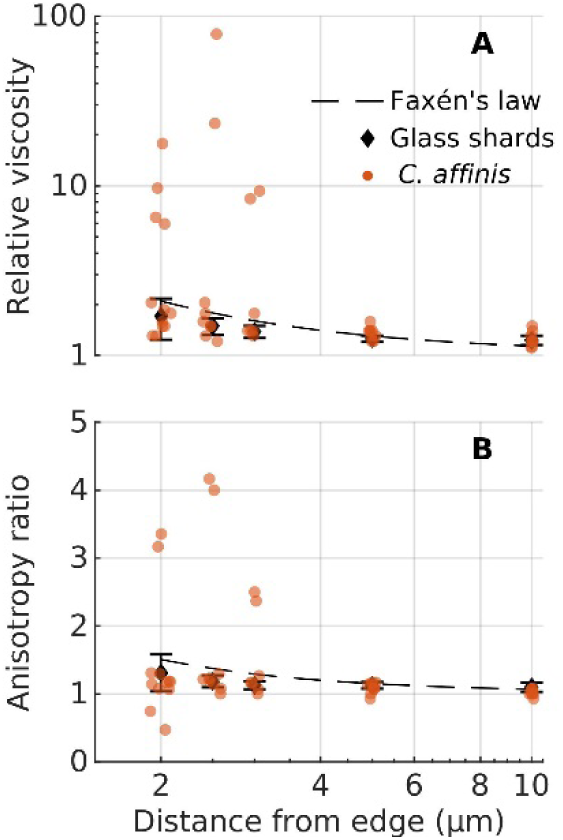
Microrheology (MOT) measurements of relative viscosity against distance from the boundary of the objects (phytoplankton cells or glass shards). Orange dots are individual measurements around *C. affinis* cells. Black diamonds are average ± st.dev. of measurements around glass shards. Dashed black lines represent predictions from Faxén’s laws (see Material and Methods). (**A**) Relative viscosity in the direction perpendicular to the edge of the object. (**B**) Anisotropy in the measurements, given as the ratio of relative viscosities in the perpendicular direction to those in the parallel direction. A value of one represents isotropic conditions. Data has been jittered in the x axis by 0.1µm to facilitate visualization.

We next used MPTM to corroborate the MOT observations, maximize the spatial coverage of the rheological measurements and investigate different species and scenarios. Maps of relative viscosity were generated around phytoplankton cells and within aggregates, with a field of view (FoV) of 130×175 µm and a spatial resolution of 2 µm. All maps can be found in (*13*). The MPTM results for healthy *C. affinis* cells (Fig. 2) are in good agreement with those obtained from MOT. In 30 of the 41 *C. affinis* cells analysed, relative viscosities near the cell wall were statistically significantly higher than those predicted by Faxén’s laws. However, we did not observe any statistically significant increases in viscosity around glass shards (Figs. 2, 3, S4 and S5, *13*), confirming that the high values recorded near the cells are not caused by hydrodynamic wall effects alone. As with MOT, relative viscosity values tend to decay nonlinearly away from the cell walls, suggesting exudation, diffusion and clustering of EPS from the cell. In all explored cases, relative viscosities measured around *C. affinis* cells were highly variable, with areas showing enhanced values and others showing no notable increase. The extent of the viscous gradient around cells was also variable, ranging from 2 to 18 µm (Fig. 3).

**Fig. 2.**
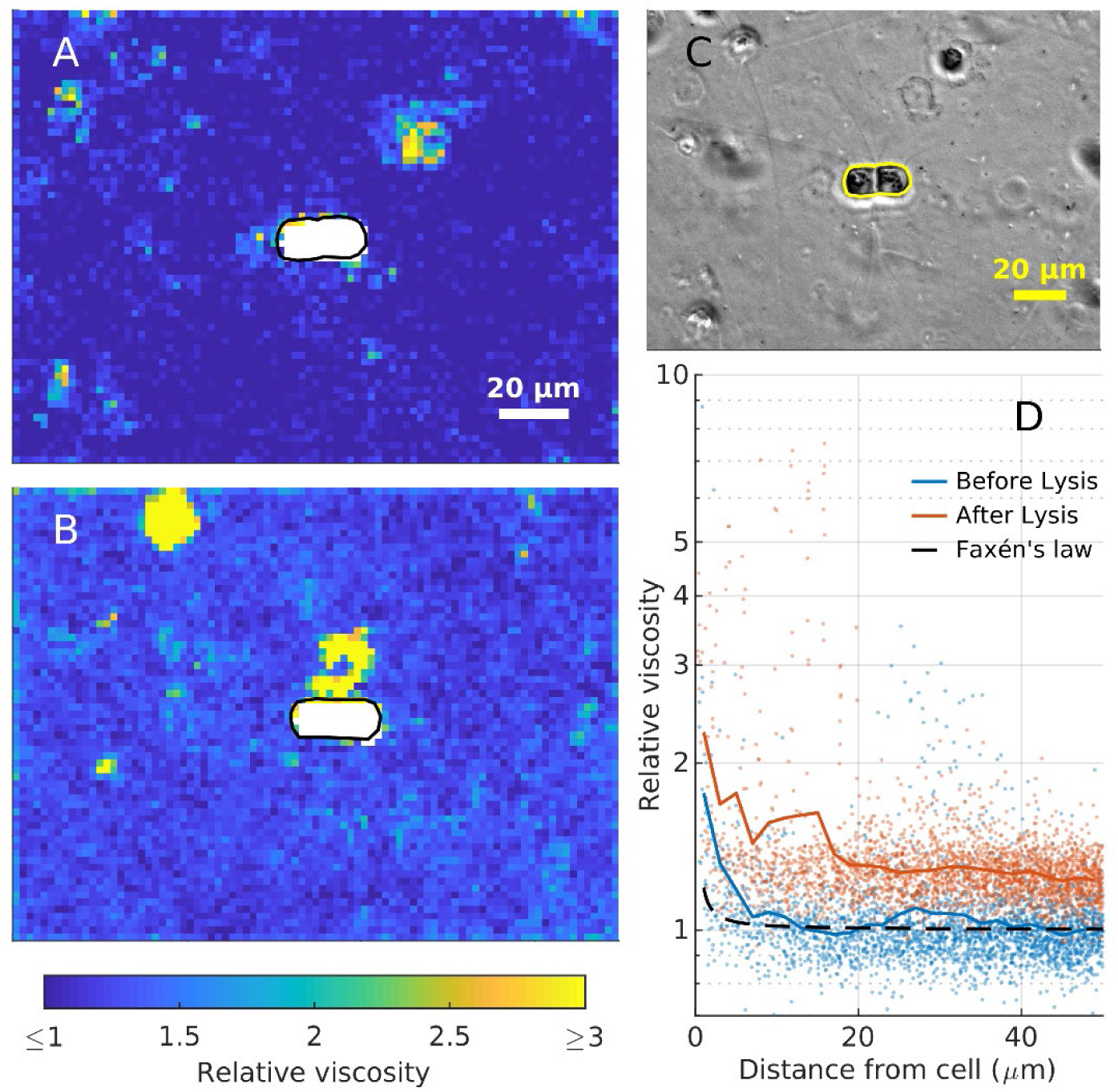
Viscosity changes around a *C. affinis* cell (**A**) before and (**B**) one hour after light-induced cell lysis. **A, B:** 2×2 µm MPTM viscosity maps. **C:** phase-contrast image at the start of the experiment. Cell boundaries are drawn in black in **A** and **B**, and yellow in **C. D:** viscosity estimates against the minimum distance to the boundary of the cell before (blue) and after (red) the lysis. Coloured lines are a moving average with a 2µm window. Dashed black line represents the Faxén’s law for motion perpendicular to a solid boundary (Eq. S2).

**Fig. 3.**
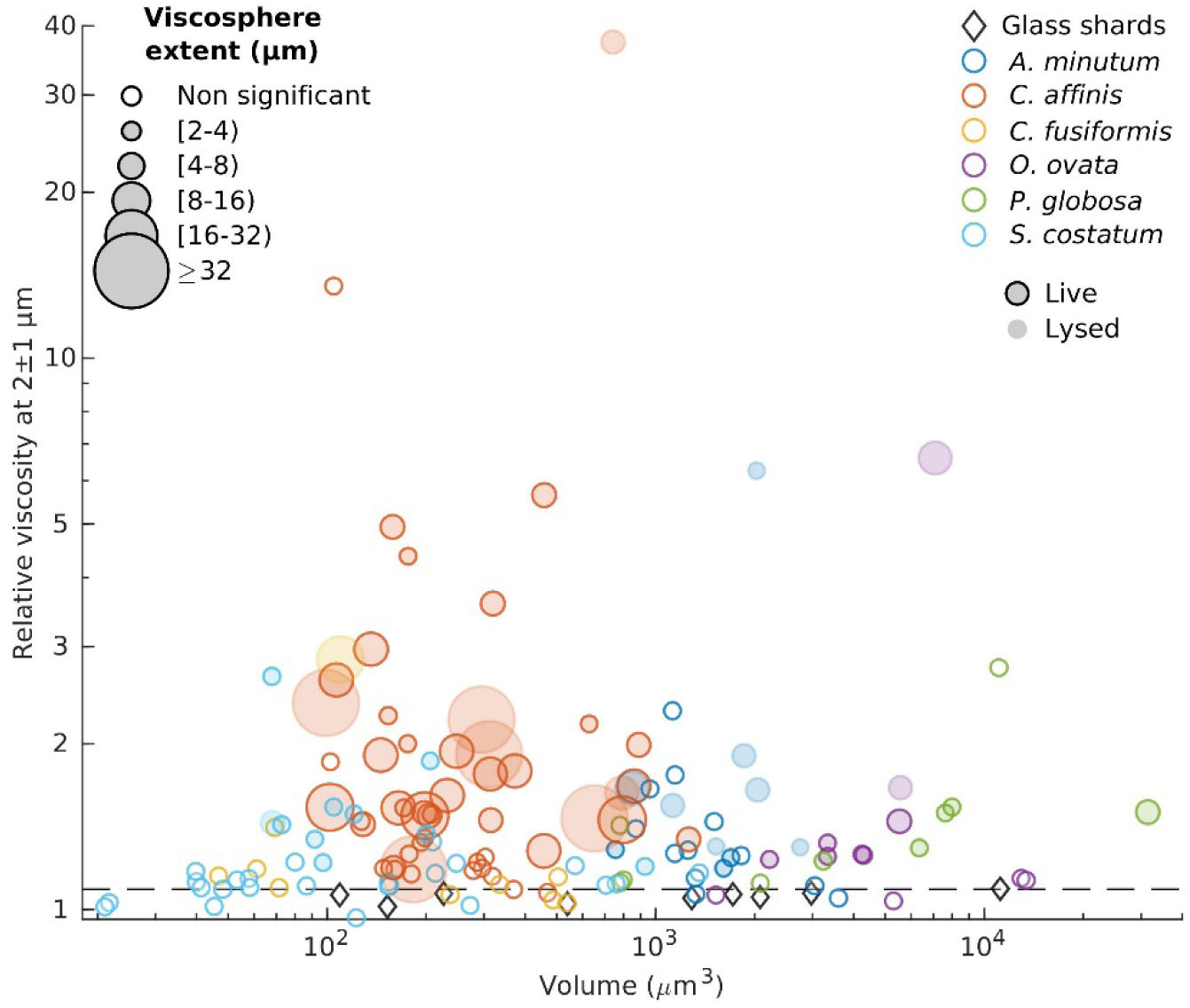
Average relative viscosities between 1 and 3 µm away from cells in relation to the cells’ volume. The diameter of each symbol is proportional to the extent of the viscous phycosphere. Horizontal dashed line represents predictions from Faxén’s law for motion perpendicular to a solid boundary (Eq. S2).

To assess the prevalence of these viscous phycospheres, which we call ‘viscospheres’, we next performed MPTM experiments on the diatoms *Cylindrotheca fusiformis* and *Skeletonema pseudocostatum*, on the dinoflagellates *Alexandrium minutum* and *Ostreopsis cf. ovata*, and on colonies of *Phaeocystis globosa*, a haptophyte whose blooms are often associated with submesoscale (∼1 km) increases in bulk-phase viscosity (*14*). We detected viscosity gradients around 45% of all cells analysed, and in all species except for *C. fusiformis* (Figs. 3, S4). The presence of a viscous phycosphere or its spatial extent, contrary to our expectations based on diffusive rates (*6*), did not correlate with cell size, while there was a high degree of intra- and interspecific variability. The spatial extent of the viscosphere for the entire dataset appeared to be exponentially distributed (Fig. S4), with large viscospheres being less frequent than small ones. Coefficients of variance of the 2D viscosity maps ranged from 8 to >2000 in all MPTM experiments (Table S1), suggesting that viscous patches are not only associated to cells and can also mark the presence of colloids and mucus sheets (*1*). The highest relative viscosity values we recorded (up to 80) were found inside the mucopolysaccharide matrix of *P. globosa* colonies (Fig S3).

The existence of a viscosphere around phytoplankton cells has profound implications beyond the physiological and ecological roles so far attributed to EPS (*15, 16*) as it fundamentally reshapes the cells’ physical environment. Exopolymers allow microalgae to structure the phycosphere and modify flow fields (*17*). Critically, an increase in viscosity reduces the rates of molecular diffusion, affecting nutrient uptake, waste removal and exchange of infochemicals. It may also influence bacterial distributions around microalgae, not only because lower molecular diffusivities lead to steeper, longer-lived and easier to track gradients of chemoattractants, but also because viscosity affects the swimming speed of bacteria (*18*) and therefore their distribution (*19*). Thus, we propose EPS exudation as an active strategy to recruit symbiotic bacteria to the phycosphere.

Viscosity gradients are also expected to develop when cells lyse, as intracellular viscosities can be several orders of magnitude higher than water (*5*). To test this hypothesis, we used MPTM to map viscosity around *C. affinis* cells before and after light-induced lysis (*20*), as well as around dead *S. pseudocostatum, A. minutum* and *O. ovata* cells. We detected increases in background viscosities (i.e. far from the cells) after exposure to UV and bright light in some *C. affinis* samples, most likely because all cells within the FoV were lysed. Additionally, we observed localized and persistent patches of high viscosity around lysed cells (Fig. 2), as well as steep viscosity gradients around dead cells. Lysing and dead cells are a hotspot of microbial activity, as chemotactic bacteria swarm around them (*20*). Therefore, we expect the same mechanisms at play in a viscous phycosphere to be magnified in a lysing cell, leading to a more efficient use of resources by chemotactic bacteria.

The last scenario we explored was of viscosity gradients generated by aggregates of phytoplankton. The formation of aggregates, TEP and marine snow is facilitated by EPS acting as a loose adhesive (*7*). We used MPTM to map relative viscosities inside and around shear-induced aggregates of the three diatom cultures. We observed increases in viscosity of more than one order of magnitude inside the aggregates (Fig. 4), with areas of enhanced viscosity largely overlapping areas stained by Calcofluor white (specific for β-d-glucopyranose polysaccharides). We regard these estimates as conservative because the spatial coverage of the MPTM maps inside the aggregates is limited by the capacity of the microspheres to penetrate the aggregates and by self-shading. Nonetheless, given the important role that aggregates play as hotspots for zooplankton foraging and microbial activity and the importance of diffusive processes in sinking of porous particles in stratified water columns (*21*), our results could pave the route to new insights into processes influencing the biological carbon pump.

**Fig. 4.**
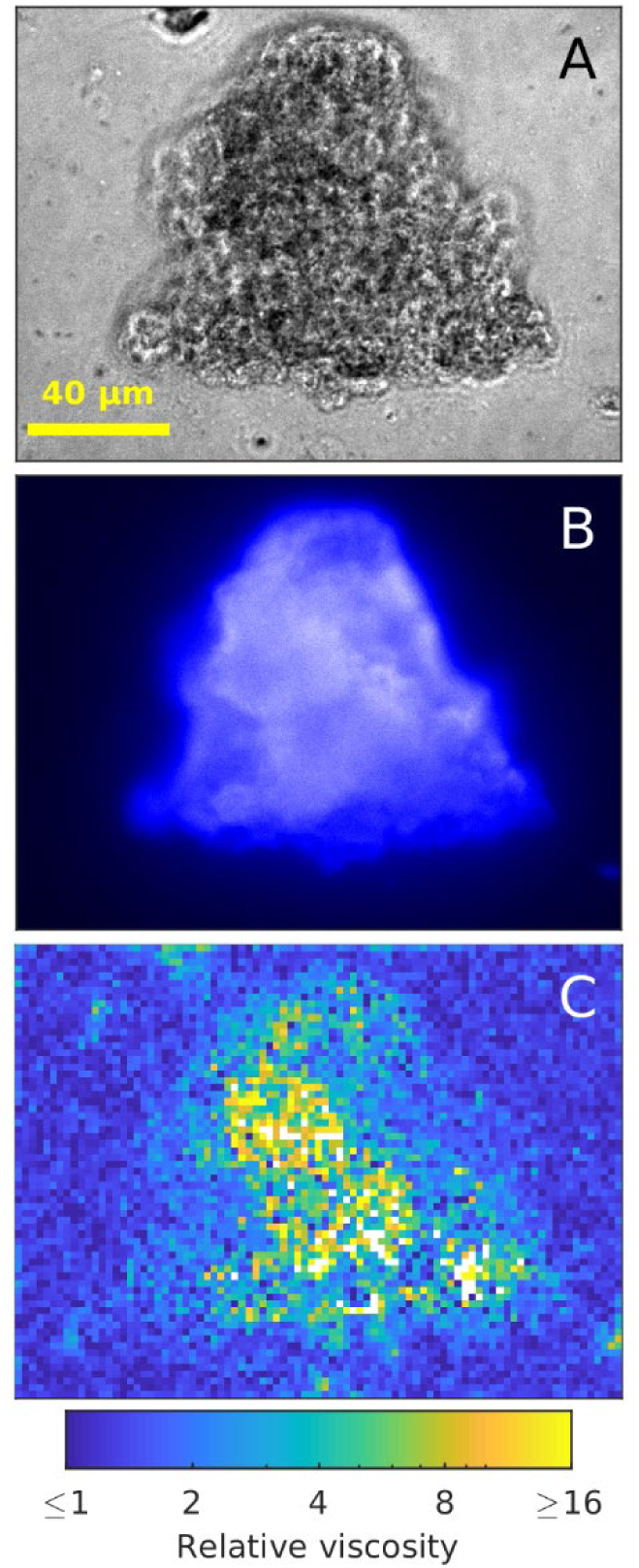
Viscosity measurements inside and around a *C. affinis* aggregate. **A**. Phase contrast image. **B** CFW staining for polysaccharides. **C** viscosity map with 2×2 µm binning obtained with MPTM. White pixels are areas with too few microsphere tracks to provide a reliable estimate.

The implications of microscale viscosity gradients are grounded in fundamental physics and are pervasive, as viscosity impacts virtually all processes and interactions occurring in the microbial world. Increasing viscosity decreases the diffusivity of dissolved substances, small passive particles and organisms, and it slows motile plankton and sinking particles. Therefore, changes in viscosity not only affect the distribution of organisms and their resources, but also slow down ecological rates, resulting in a cascade of effects at different ecological levels and scales. Increasing viscosity around osmotrophs decreases nutrient uptake rates, impacting primary and bacterial productivities. Similar reductions in encounter rates between predators (or viruses) and prey (or hosts) along with uncertain effects on the formation and sedimentation of aggregates, ultimately influence the transfer of carbon across trophic levels and the strength of the biological carbon pump. Finally, given that viscosity decreases nonlinearly (e.g. exponentially for simple fluids) with increasing temperature, we must expect changes in viscosity in the context of global climate change, with the potential for unforeseen consequences for the carbon cycle and climate regulation (*22*). In summary, the inclusion of viscosity adds a new layer of complexity to the current paradigm of microscale heterogeneity and our results provide a quantitative framework to explore how these ideas unfold from the microscale upwards.

## Acknowledgements

We thank E. Berdalet, F. el Baidouri, R. Holzman, C. Marrasé, F. Peters, R. Schuech and R. Simó for insightful comments on previous versions of this manuscript. We thank E. Berdalet for providing *O. ovata* and S. Vaidyanathan for providing *C. Fusiformis*.

## Author contributions

O.G. and S.H. designed the study. A.J.W., M.T. and T.M. developed the methodology and T.M. performed the experiments for MOT. M.T. developed the software for MOT analysis. O.G. developed the methodology and software for MPTM. O.G. and M.S.-N. performed the MPTM experiments. A.J.W. and M.T. provided guidance on the use and significance of microrheology. O.G and T.M. wrote the first draft, and all the authors discussed the results and contributed to the writing of the manuscript.

## Data and materials availability

All MPTM viscosity maps have been deposited in figshare (*13*), as well as the code used for MPTM experiments and analysis (*23*). Any additional data that support the findings of this study are available from the corresponding author upon request.

## Materials and Methods

*Phytoplankton species and cultures. Formation of aggregates*.

### Growth conditions

We performed our experiments on the diatoms *Chaetoceros affinis* (CCAP 1010/27), *Cylindrotheca fusiformis* (CCAP 1017/2) and *Skeletonema pseudocostatum* (CCAP 1077/7), the dinoflagellates *Alexandrium minutum* (CCAP 1119/15) and *Ostreopsis cf. ovata*. (OOPM17), and the haptophyte *Phaeocystis globosa* (strains K-1321 and K-1323, Norwegian Culture Collection of Algae). Diatoms were grown in f/2 + Si medium (*24*) in 33.5g/L artificial sea water (ASW, Aquarium Systems Instant Ocean Salt). *A. minutum* and *P. globosa* K-1323 were grown on L1 medium (*25*) in 30g/L ASW. *P. globosa* K-1321 was grown on TL medium (Norwegian Culture Collection of Algae) in 30g/L ASW. All species, except *O. ovata*, were grown at 19 degrees Celsius without shaking in an algae incubator (Algaetron AG 230, Photon Systems Instruments, Drasov, Czech Republic) with a 12:12h illumination cycle. The light intensity was 100µE for diatoms and 90 µE for *A. minutum* and *P. globosa. O. ovata* was grown with L1 medium in 38.5 g/L ASW and kept at lab temperature (20°C) at the windowsill to ensure it received enough illumination. *C. affinis* cultures for the MOT experiments were also grown at room temperature on a windowsill. Since cultures were not axenic, to ensure few bacteria were present during the experiments, these were carried out less than two weeks after culture refreshment.

Aggregates of cells were formed from *C. affinis, C. fusiformis* and *S. pseudocostatum* cultures by incubating newly refreshed monocultures on a roller table (*26*). The roller was on the windowsill with natural light conditions at 20°C. Within a few hours, aggregates were visible. Individual aggregates were collected very gently with a capillary tube to minimize disruption and prepared for MPTM analysis using the same protocol than for phytoplankton below.

#### Optical trapping microrheology (MOT)

Microrheology with optical tweezers (MOT) uses single, optically trapped, microspheres as probes for measuring the rheological properties of the material surrounding them. Each microsphere is held by a tightly focused laser beam and can be manipulated in 3D space, allowing it to be placed at specific locations around an object of interest (e.g. a phytoplankton cell).

A microsphere confined within an optical trap will move with amplitudes on the nanometre scale due to the Brownian motion of the molecules of the surrounding material. The time dependent mean-squared displacement (MSD) calculated from the residual motion of a trapped microsphere can be used to compute the rheological properties of the fluid around it as well as the strength of the optical trap (*27-30*). For sufficiently long measurements the relative viscosity (defined as the ratio of measured viscosity to that of ASW) can be calculated by plotting the particle normalised position autocorrelation function (NPAF) against time (*29*).

In this study MOT was performed with polystyrene microspheres (diameter = 2.12 µm; 19508-2, Polysciences, Inc.) as optically trapped probes for measuring the viscosity of the ASW (f/2 + Si medium) at specific positions around individual *C. affinis* cells. The phytoplankton culture was refreshed the day before each experiment. The microspheres were dispersed at a final concentration of 0.0001% v/v in a dilute (10% v/v) solution of the *C. affinis* culture in f/2 + Si. This solution was loaded into a µ-slide 8 well glass-bottom coverslips (ibidi GmbH) and incubated for ∼2 hours at room temperature. The coverslips were pre-coated with poly-L-lysine (Sigma Aldrich P4707) to immobilize the phytoplankton cells and to prevent them from being drawn into the optical trap. Control samples were prepared in the same coated µ-slide coverslips using inert glass shards (from crushed coverslips – Borosilicate glass, 0.17µm thickness; Thermo Fisher Scientific) suspended in f/2 + Si medium.

The microspheres were trapped using a 1064 nm continuous wave laser (Ventus 1064, Laser Quantum Ltd., UK) with a laser power of <5mW at the sample, and a high numerical aperture (NA) objective lens (100x 1.3NA Nikon CFI Plan Fluor). The trap depth was kept constant at 10µm from the coverslip throughout the measurements while the lateral position was varied. Each measurement involved a different microsphere and phytoplankton cell due to the challenges of retaining the same microsphere and cell for each sequence of positions. Each trapped microsphere was imaged in wide-field transmission on an inverted microscope (Eclipse Ti, Nikon) using a fast CCD camera (DALSA Genie, Teledyne Technologies International Corp). The time-dependent MSD of the trapped microspheres were tracked in real time and in 2D across 200,000 frames recorded at a frame rate of 1 kHz using a bespoke LabVIEW (version 2013 32bit, National Instruments) VI (virtual instrument) (*31*). The relative viscosity of the media around the cells/glass shards was calculated from the tracked coordinates using a previously published LabVIEW VI (the executable is free to download together with the instructions from the following link: https://sites.google.com/site/manliotassieri/labview-codes). Care was taken to ensure that the angle of approach of the trapped bead to the edge of the glass/cell was 90° and when this was not possible the tracked position data was rotated by the offset angle as part of the analysis.

### Edge effects

A moving sphere experiences an increase in hydrodynamic drag at close proximity to a solid surface. Since the hydrodynamic drag coefficient *γ* is proportional to the dynamic viscosity *η* of the fluid, this effect translates into an apparent increase in the viscosity of the fluid near the boundary. This increase in *γ* has been estimated by Faxén’s laws (*32, 33*) and depends on the radius of the sphere *a*, its distance from the surface *s*, and the viscosity of the material in which the sphere is suspended. Faxén’s effect experienced by a microsphere moving parallel to the surface is given by (*34*):

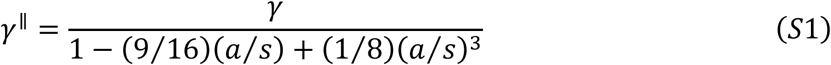

where *γ*^║^ is the corrected hydrodynamic drag coefficient for a sphere moving parallel to the surface, *γ* is the hydrodynamic drag coefficient far from the surface, which for a sphere is *γ=6* π *a η*. Similarly, for the microsphere’s motion perpendicular to the surface the Faxén effect is:

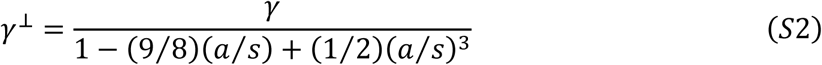

where *γ*^┴^ is the corrected hydrodynamic drag coefficient for a sphere moving perpendicular to the surface. The apparent increase in relative viscosity due purely to hydrodynamic effects close to a boundary wall, in the parallel and perpendicular directions, is given by the ratio of the drag coefficients *γ*^║^/ *γ* and *γ*^┴^/ *γ*, respectively.

### Anisotropy

We assessed the anisotropy of the local relative viscosity by calculating the ratio of the relative viscosity measured from the particle motion perpendicular to the edge of the cell or glass shard by the relative viscosity measured from the particle motion parallel to the edge of the cell or glass shard. Figure 1B includes Faxén’s effect and hence the effects purely due to the hydrodynamics near a solid wall by plotting gamma perpendicular divided by gamma parallel for varying distances from the edge of the object.

#### Passive multiple particle tracking microrheology (MPTM)

To assess spatial variability in viscosity at the microscale, we used passive multiple particle tracking microrheology (*35*). Briefly, the sample of interest is seeded with fluorescent microspheres of known diameter and density. The microspheres are tracked under the microscope as they undergo free Brownian motion. From the analyses of their trajectories, and given a known and constant temperature, it is possible to estimate the local dynamic viscosity of the fluid into which the microspheres are suspended. We partitioned the 2D field of view into squares of given length and estimated the average dynamic viscosity from the analysis of all the tracks registered within each square. This procedure allowed us to generate maps of relative viscosity with a resolution of 2µm, around individual phytoplankton cells as well as around, and to certain extent within, aggregates. All our maps have been published without any preselection in a figshare collection (*13*).

### Sample preparation and visualization

To measure local relative viscosity, a 1 mL sample of the culture of interest was seeded with 50 µL of a solution of fluorescent polystyrene beads of 0.29 µm diameter (Fluoresbrite® YG Carboxylate Microspheres, Polysciences Inc.). The solution of microspheres was prepared by diluting one drop (∼40 µL) of the commercial solution into 2 mL of the ASW used to prepare the microalgae media. A surfactant (Tween 20), was added to the solution at 0.3% to facilitate the disaggregation of microspheres. Then, the microspheres solution was mixed on a vortex and sonicated in a bath sonicator for 5 minutes. The density of the microspheres is 1.05 g·mL^-1^ compared to 1.024 g·mL^-1^ of ASW at 33.5 salinity at 20°C. Under these conditions the microspheres would sediment at ∼1nm·s^-1^ (*35*). The final concentration of microspheres in the solution was 2·10^9^ microspheres m·L^-1^ and the volume fraction was 2·10^−5^. From these values it is possible to calculate an average distance between microspheres of 9 µm (*35*).

Specimens were prepared by pipetting 100-150µL of the sample with microspheres on a slide in between two coverslips. The sample was then topped with a coverslip, sealed with vaseline to suppress any drift, and placed on an inverted epifluorescence microscope (Zeiss AxioVert A1 Fl) equipped with an LED source (CoolLED pE-300 white) and imaged at 400X magnification with a long-distance objective (ZeissLD Plan-Neofluor 40x/0.6NA Corr). Pictures and videos were taken with a sCMOS camera (Hamamatsu ORCA Flash 2.0). The field of view was 131 by 175 µm and the depth of field was 1.45 µm. To perform an analysis of the relative viscosity, we selected glass shards, cells or aggregates that were isolated and completely still. Once the shard, cell or aggregate was selected, the fluorescent illumination was switched on and several consecutive videos were taken at 30 frames per second. Intensity of light was set to a maximum of 5% to avoid heating the sample or eliciting any response by the phytoplankton or bacteria. Note that the analyzed *O. ovata* cells showed a round shape that could correspond to cysts (*36, 37*). Cyst formation can be rapidly triggered by the stress/shear associated to manipulation procedures.

We have identified the following potential sources of error in our method: inaccuracy of the temperature control, mechanical vibrations in the sample, inclusion of the analyses of tracks of clusters of microspheres and imprecision in the detection of particles and the estimation of their centroids. The first two error sources were minimized by controlling temperature throughout the experiments with a temperature stage (PE100-ZAL System, Linkam Scientific Instruments Ltd., Tadworth, UK) and by placing the microscope on an active antivibration table (ScienceDesk, Thorlabs Inc. Newton, New Jersey, United States). Furthermore, microsphere aggregates and other particles larger than single microspheres were discarded by the particle detection code.

When developing this methodology, we observed that microspheres could occasionally attach to cells or aggregates and move together with them. When analysing the tracks from these microspheres we obtained estimates for viscosity that could be several orders of magnitude higher than that of water because they were diffusing at the same rate than the larger particle. To avoid this potential source of error, we used Polysine™ slides (Thermo Scientific™) and targeted only cells, aggregates and glass shards that had attached to the slide and were completely still. To ensure this we took phase-contrast pictures before and after the tracking and discarded any videos in which the cells or particles of interest had moved. We focused ∼10 µm above the slide to avoid wall effects and to image only microspheres that were freely moving. Preliminary tests with the Polysine slides confirmed that any wall effects were below the precision of the method (see below) and that they were adequately accounted by Faxén’s laws (Fig S5). As with MOT, control samples with inert glass shards (from crushed coverslips – Borosilicate glass, 0.17 µm thickness; Thermo Fisher Scientific) were prepared on Polysine slides.

### Lysi

of individual phytoplankton cells was induced by exposing them to UV and white light at maximum intensity for approximately 10 minutes, following Smriga et al. (*20*). The field of illumination was narrowed to minimize effects on nearby cells. In the case of diatoms, the lysing procedure was deemed successful when motile bacteria started to aggregate around the cell (signalling release of material). To minimize hydrodynamic interactions of swimming bacteria with the microspheres, we waited until the cloud of bacteria had dissipated before starting the recordings, which could take at least 10 minutes but no more than an hour. In the case of dinoflagellates, which are motile, we immobilized the cells by exposing them to UV and white light at the maximum intensity for a short period of time. Upon exposure to intense light *A. minutum* cells would first loose the lateral flagellum and then the polar one. If the light was left on for longer, a ‘blob’ of material would be seen coming out of the cell.

### Viscosity calculation

The field of view was partitioned into 2 by 2 µm squares. For each individual microsphere trajectory, we computed the squared displacements with lag-times *τ* between 1/30 and 5/30 seconds. The squared displacements were grouped into the partitions according to their mean centroid, and the ensemble mean square displacements (*MSD*s) were computed for each partition. The diffusivity *D* of the microspheres within each partition was calculated by least squares fitting the data to:

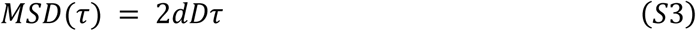

where *d* = 2 is the dimensionality of the tracks. The average dynamic viscosity *η* for each position was then calculated from the Stokes-Einstein equation for diffusion of a spherical particle:

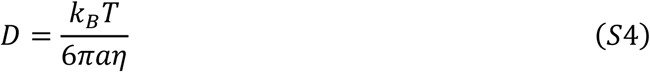

where *k*_*B*_ is the Boltzmann constant and *T* is the absolute temperature.

Background dynamic viscosities were calculated as the median of measured dynamic viscosities for the entire field of view. Relative viscosities were calculated dividing the local dynamic viscosities by the dynamic viscosity of ASW estimated using equations in (*38*).

The image acquisition and analysis were conducted with in-house Matlab code. The microsphere tracking used a Matlab code initially designed to track bacterial cells (*39*). All the code, including the code used to estimate local relative viscosities and create the maps, is available online under an GPL 3.0 license (*23*).

### Viscosity profiles

To characterize the decay in viscosity away from a particle, the boundary of the particle was first characterized manually on the image using the command *drawpolygon* (*drawcircle* in the case of spherical *P. globosa* colonies) from the Image Processing Toolbox in Matlab. For each 2×2µm partition, the distance to the particle was computed as the minimum Euclidean distance to any of the boundary vertices.

A conservative estimate of the minimum length of the viscosity gradient was calculated for each particle in the following way. Viscosities were grouped into 2 µm wide bins according to their minimal distance to the boundary of the particle of interest. For each group a non-parametric Wilcoxon signed rank right-sided test was conducted with a 5% significance threshold to assess whether median viscosity was higher than the value predicted by Faxén’s laws. To minimize the number of false positives, we used the Faxén’s equation for movement perpendicular to the solid edge (eq. S2) which gives the highest value. The minimum length of the viscosity gradient was then estimated as the mid value of the bin immediately smaller than the first bin at which the median viscosity was significantly lower than the Faxén’s prediction.

The volume of the cells was estimated assuming an ellipsoid shape where the two minor axes were assumed to be both equal to the width of the outlined cell boundary, while the major axis was equal to the length.

### Method calibration and uncertainty

To quantify uncertainty, we first checked whether our method was systematically biased. We performed a series of microrheological experiments on ASW with a salinity of 33.5 at a range of different temperatures, from 5°C to 35°C, and compared the results with the predictions from empirical equations for seawater (*38*). The slope of the least squares linear regression model was not significantly different from unity (Fig S6, *T*-test: *t*(7) = 1.098, *p* = 0.309), although our estimates were systematically higher than the theoretical prediction. We calculated this bias to be 2.4 10^−4^ (kg m^-1^ s^-1^). In all subsequent calculations we subtracted this bias from our estimate for the absolute dynamic viscosity.

We further tested our method outside the limited range achievable through changing temperature by comparing it against bulk measurements performed with a cone-plate rotational viscometer (DV2T Brookfield). We prepared a series of solutions of increased viscosity adding Ficoll 70, a highly stable, long chain synthetic polymer of fructose, to water. Aqueous solutions of Ficoll are Newtonian at the range of concentrations used in our study (*40*). Ficoll solutions were prepared by dissolving the polymer in ultrapure water and kept in a shaker rotating at 200RPM overnight. The results, after accounting for the bias with theoretical estimates, lie on a 1:1 slope for the entirety of the range (Fig S6).

We evaluated the precision of viscosity estimates in 2×2 µm square by calculating the bootstrap coefficient of variation (*CV*) from the seawater and Ficoll experiments outlined above. We estimated the unbiased *CV* as:

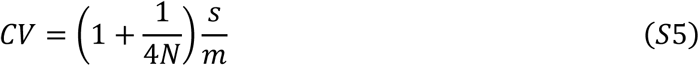

Where *N* is the number of tracks within a 2×2 µm square and *s* and *m* are the sample standard deviation and mean, respectively. The precision of viscosity estimates decreases nonlinearly with the number of tracks registered in a bin and does not depend on the viscosity (Fig. S7). In subsequent analyses, we used a minimum threshold of 20 tracks per bin to accept the viscosity estimate for any particular bin. This ensured a good coverage with a maximum *CV* of 20%. It should be noted that in most pixels the actual *CV* was much lower than 20%, as the average number of tracks was usually much higher than 20.

### Exopolymer staining

Simultaneously with the MPTM of aggregates we assayed staining of the sample with several labels commonly used to visualize different EPS components in biofilms and flocs (*41, 42*). Explicitly, we tested (*i*) Alcian blue (Sigma-Aldrich A5268), which is specific for negatively charged polysaccharides commonly used to dye transparent exopolymer particles (*43*), (*ii*) FilmTrace SYPRO Ruby (TermoFisher Scientific F10318), which labels most proteins, (*iii*) SYBR Green I, which labels nucleic acids, (*iv*) Concanavalin A-Tetramethylrhodamine conjugate (Con A, Invitrogen™ C860), which labels α-d-Glucose and α-d-mannose and (*v*) Calcofluor white M2R (CFW, Sigma-Aldrich 18909), which labels β-d-glucopyranose polysaccharides. Alcian blue revealed an intricate network of polysaccharides within our samples, but we did not use it alongside MPTM because it precipitates in the presence of salts, creating a rigid matrix that alters the rheological properties of the fluid. Of the other stains, Concanavalin A and CFW produced the highest signal. In subsequent analyses of aggregates we used CFW.

## Supplementary figures

**Fig. S1.**
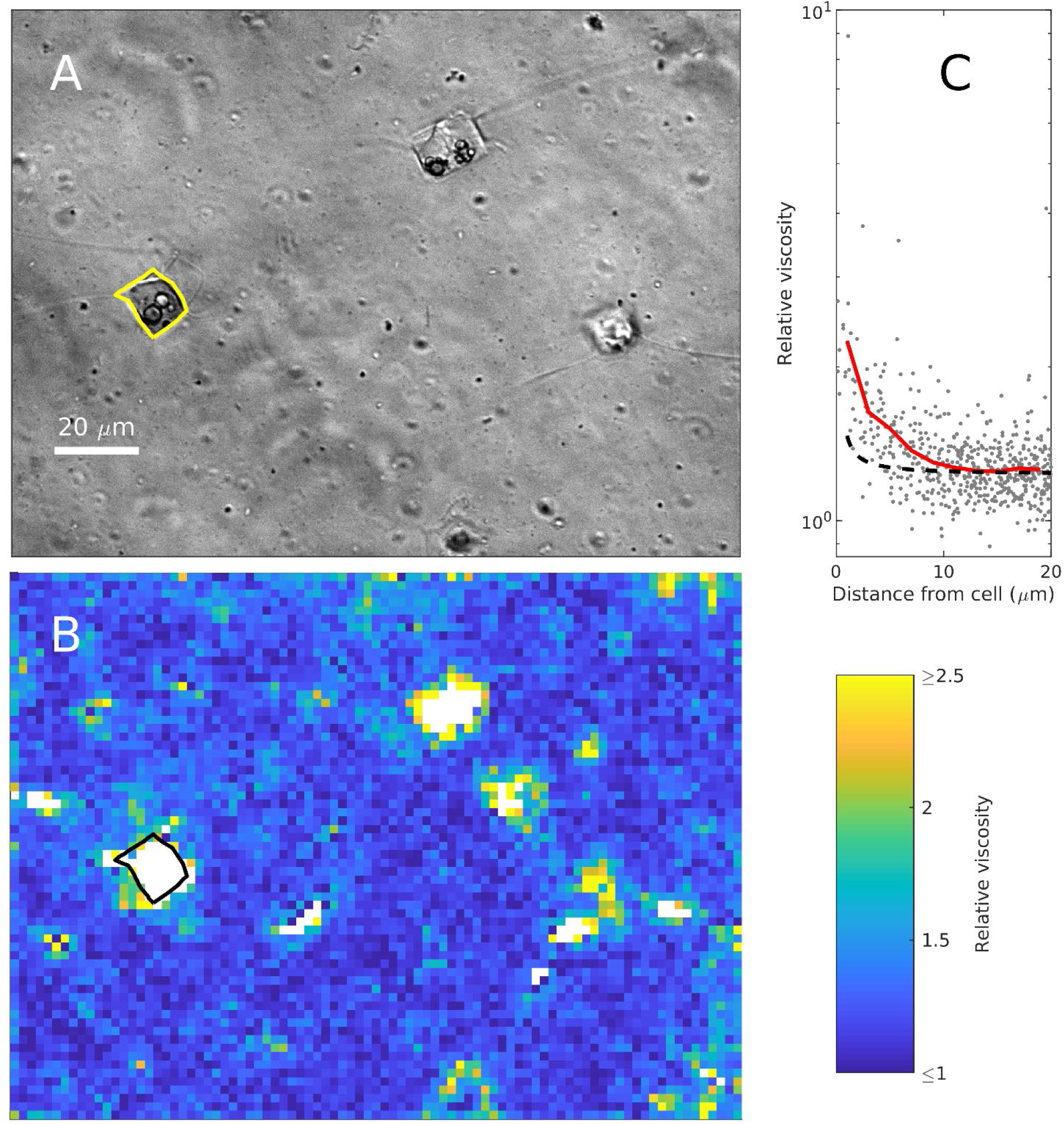
Viscosity map and average profile around a healthy *C. affinis* cell. **A**: phase contrast image at the start of the experiment. **B**: map of viscosities estimated using MPTM. **C**: viscosity estimates against the minimum distance to the boundary of the cell. Red line in **C** is the moving average with a 2µm window. Boundary of the cell analysed is overlapped in yellow in **A** and in black in **B**. Black dashed line in **B** represents predictions from Faxén’s laws for motion perpendicular to a solid boundary (Eq. S2).

**Fig. S2.**
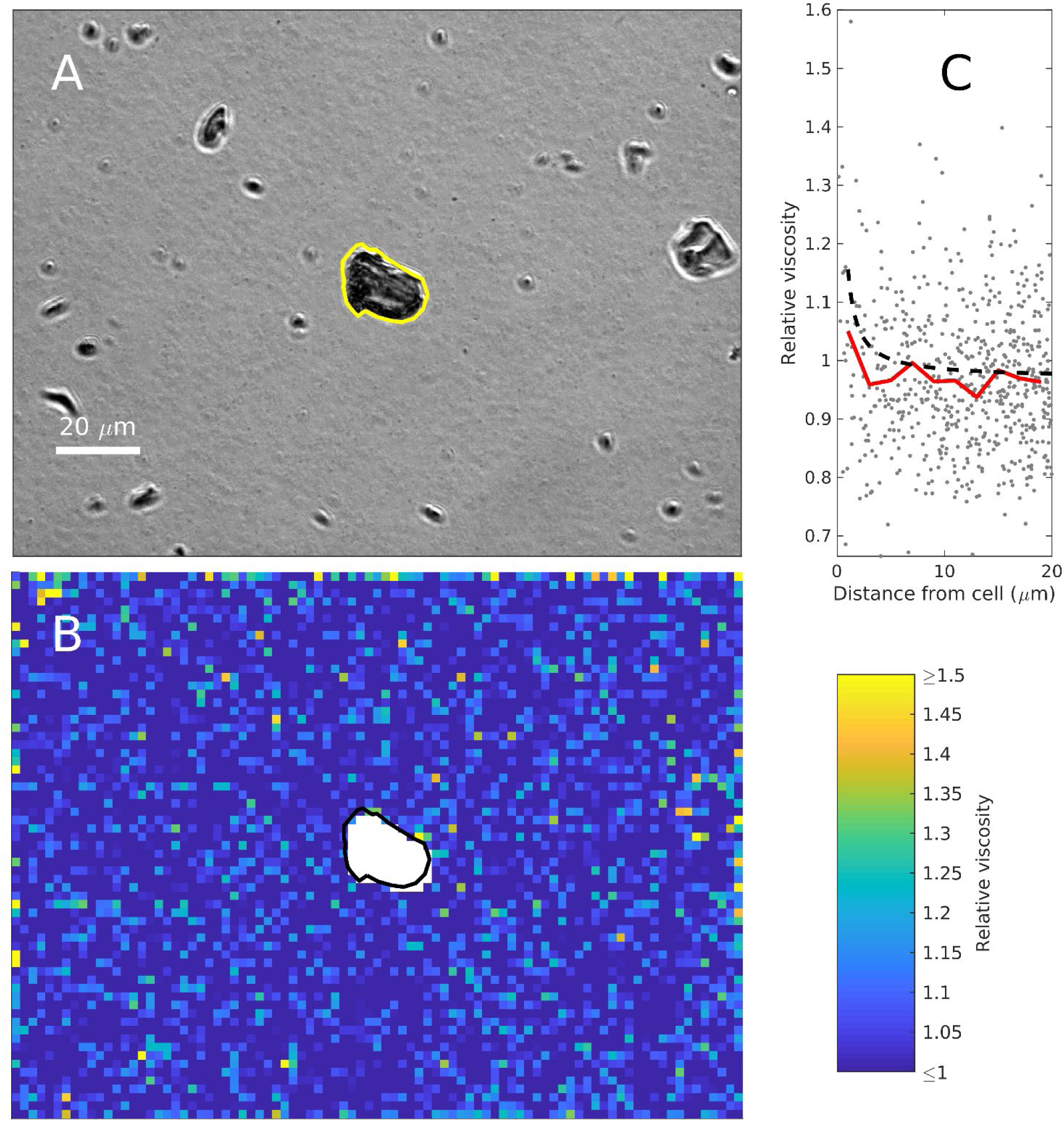
Viscosity map and average profile around a glass shard. **A**: phase contrast image at the start of the experiment. **B**: map of viscosities estimated using MPTM. **C**: viscosity estimates against the minimum distance to the boundary of the shard. Red line in **C** is the moving average with a 2µm window. Boundary of the shard analysed is overlapped in yellow in **A** and in black in **B**. Black dashed line in **B** represents predictions from Faxén’s laws for motion perpendicular to a solid boundary (Eq. S2).

**Fig. S3.**
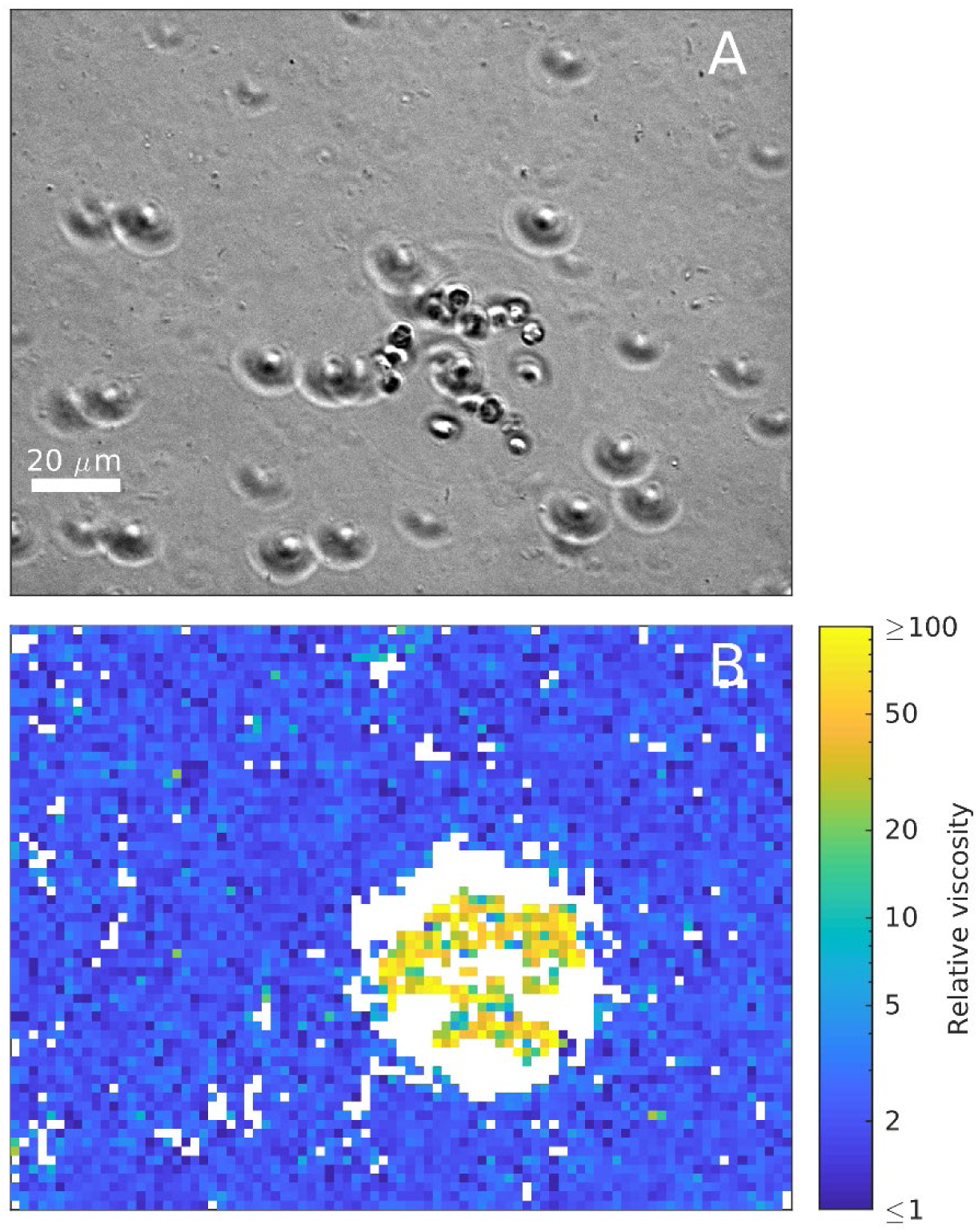
Viscosity map of a *P. globosa* colony. **A**: phase contrast image at the start of the experiment. **B:** map of viscosities estimated using MPTM. White pixels are areas with too few microsphere tracks to provide a reliable estimate.

**Fig. S4.**
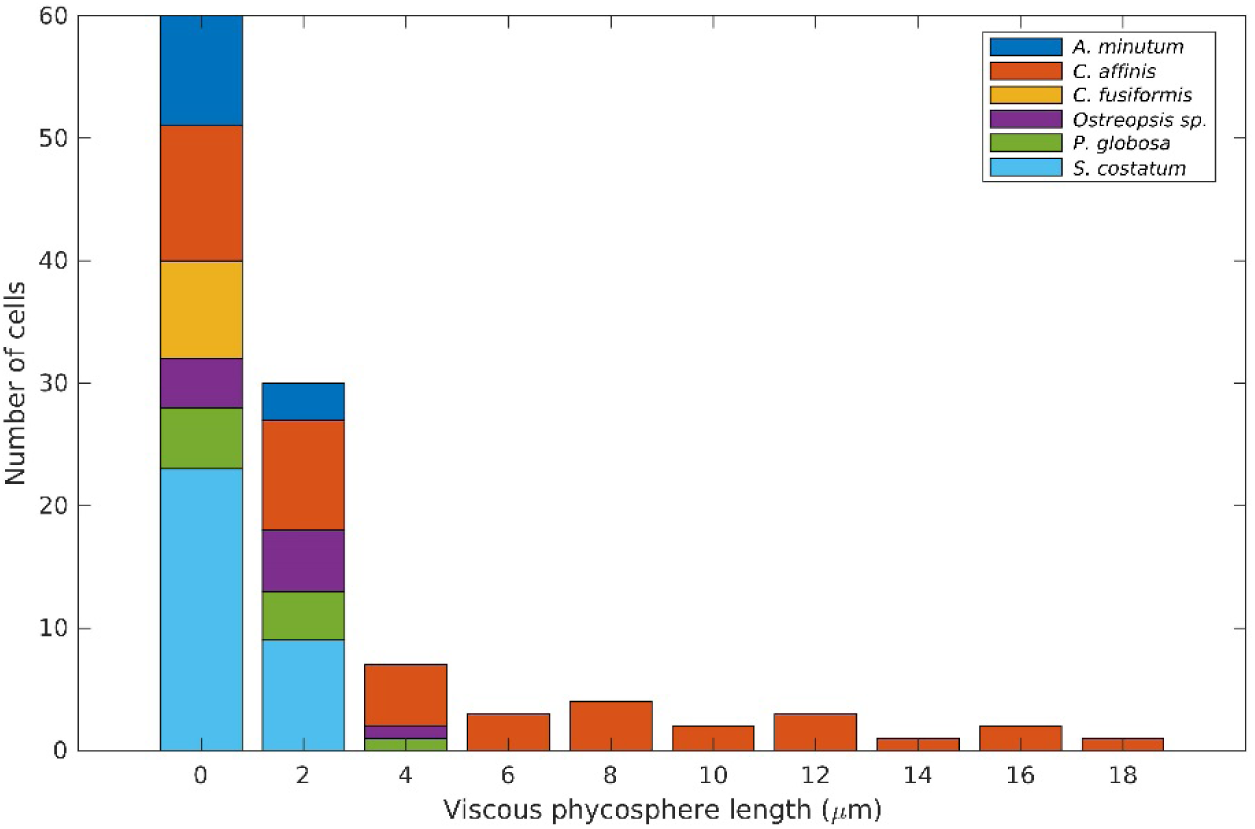
Frequency distribution of the spatial extent of the viscous phycosphere among the phytoplankton cells analysed.

**Fig. S5.**
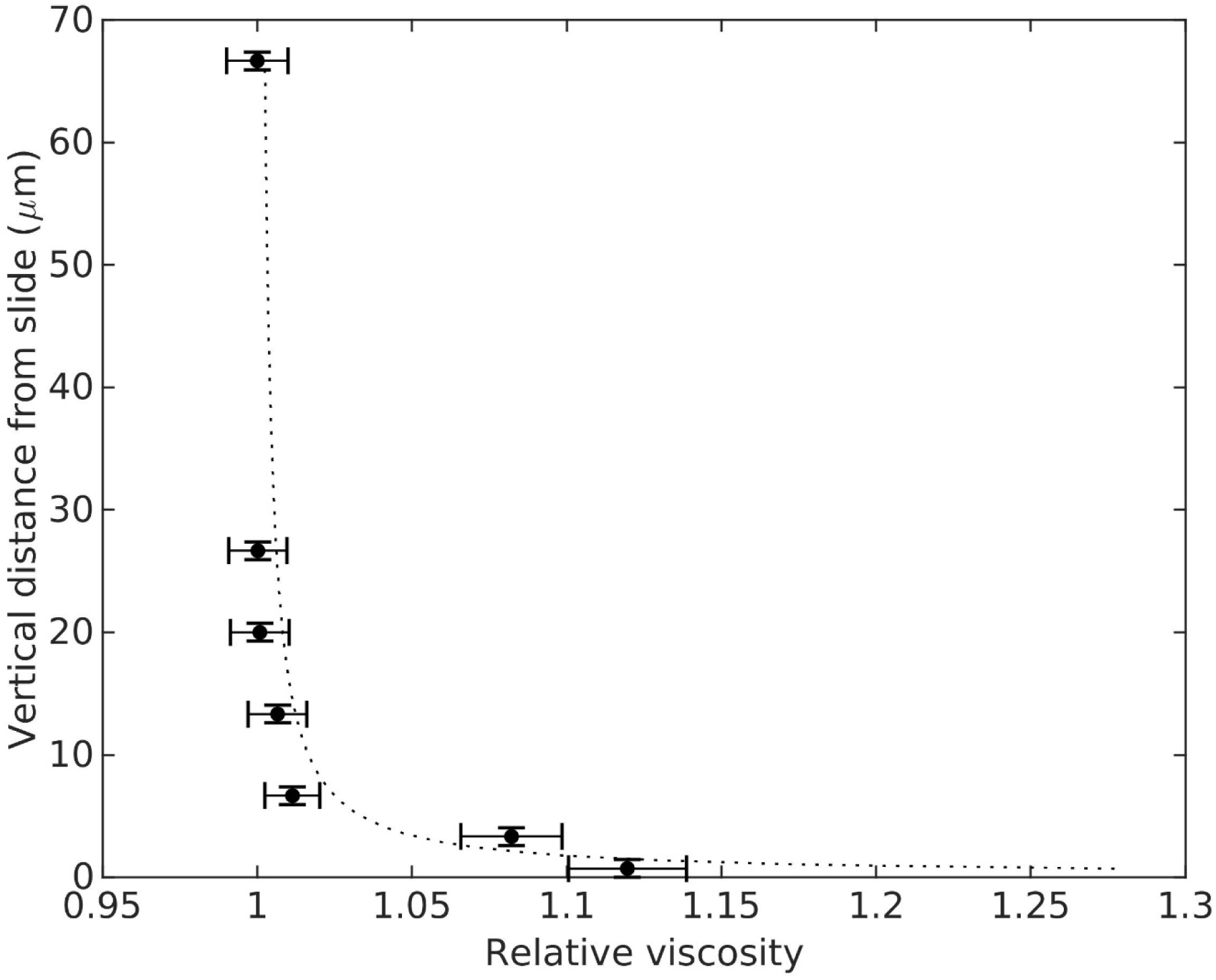
Average relative viscosity of ASW with 33.5 salinity at 21°C calculated with MPTM *vs* vertical distance from the slide. Horizontal bars are the standard errors. Vertical bars represent the depth of field. The dashed line is the apparent relative viscosity predicted by Faxén’s law (eq. S1).

**Fig. S6.**
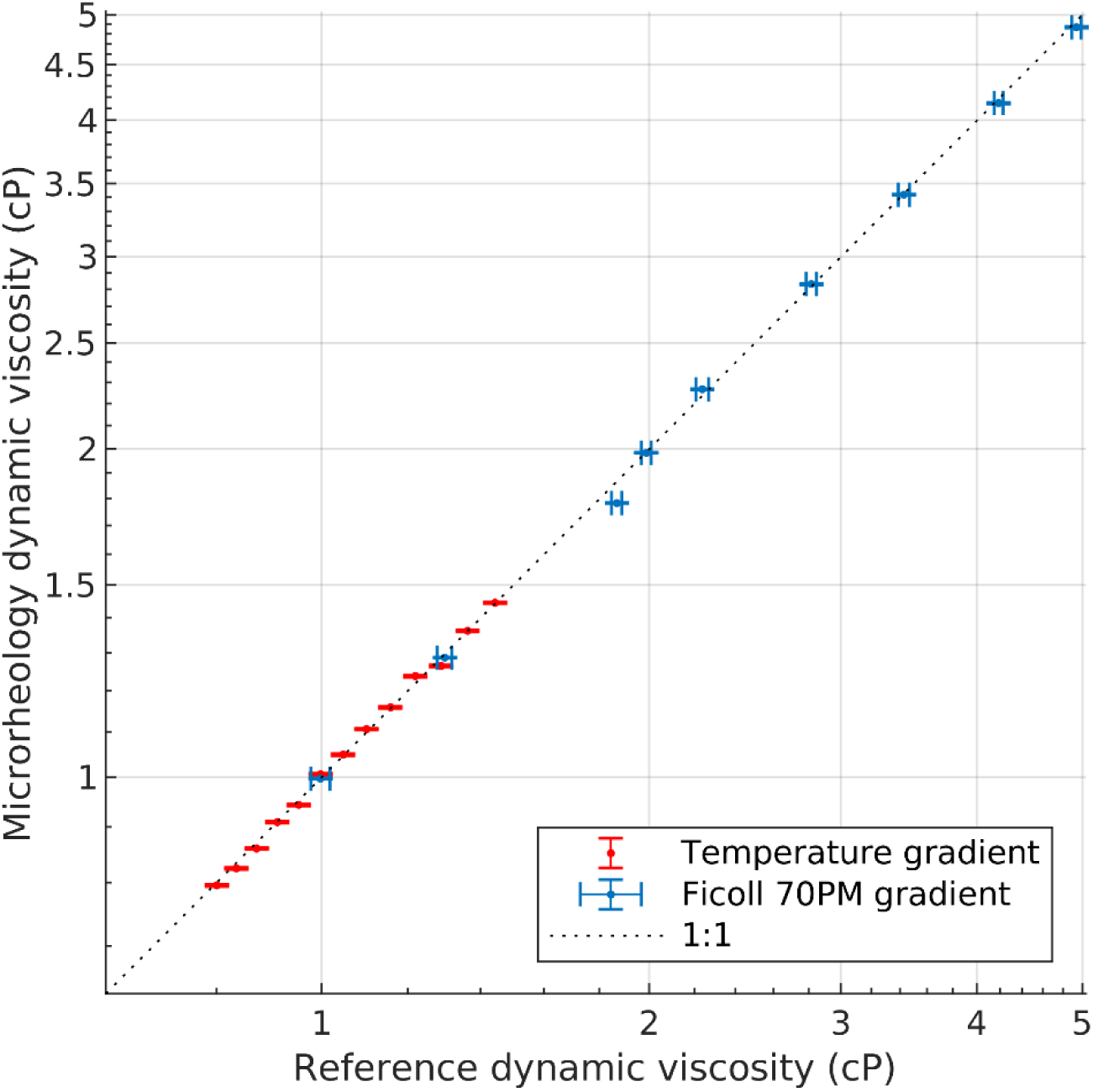
Dynamic viscosities of ASW at different temperatures (in red) and water solutions of Ficoll 70PM, estimated using MPTM against reference estimates. The reference viscosities for the temperature gradient were calculated using empirical equations in(*38*). The reference viscosity for the Ficoll gradient were measured with a DV2T Brookfield cone-plate viscometer. Vertical error bars show the standard errors. Horizontal error bars show the reported precision of the viscometer.

**Fig. S7.**
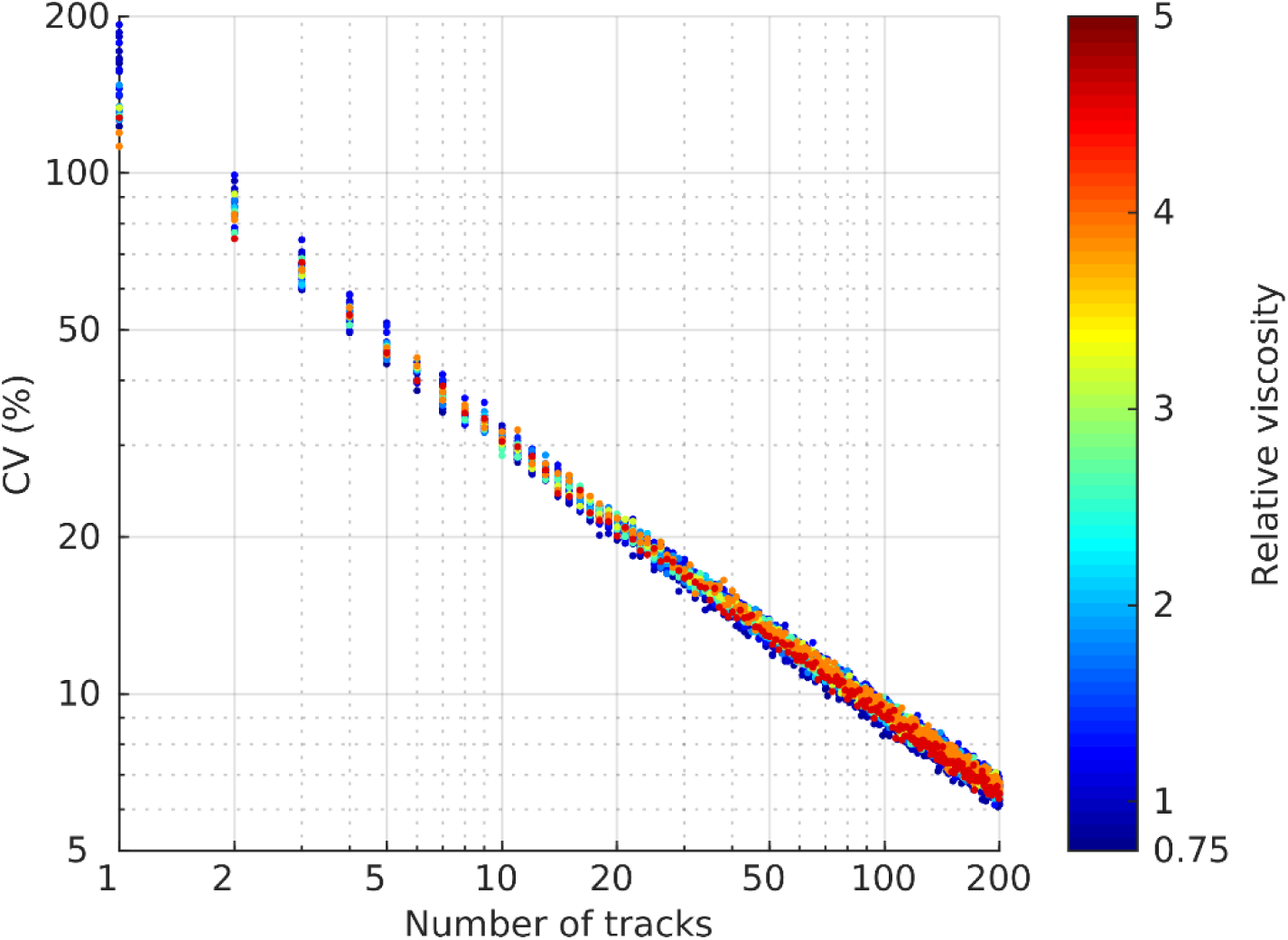
Bootstrapped coefficient of variation against number of tracks per bin, which can be considered as the error for each pixel in the viscosity maps. Colour indicates the average viscosity of the sample used.

## Supplementary table

**Table S1.**
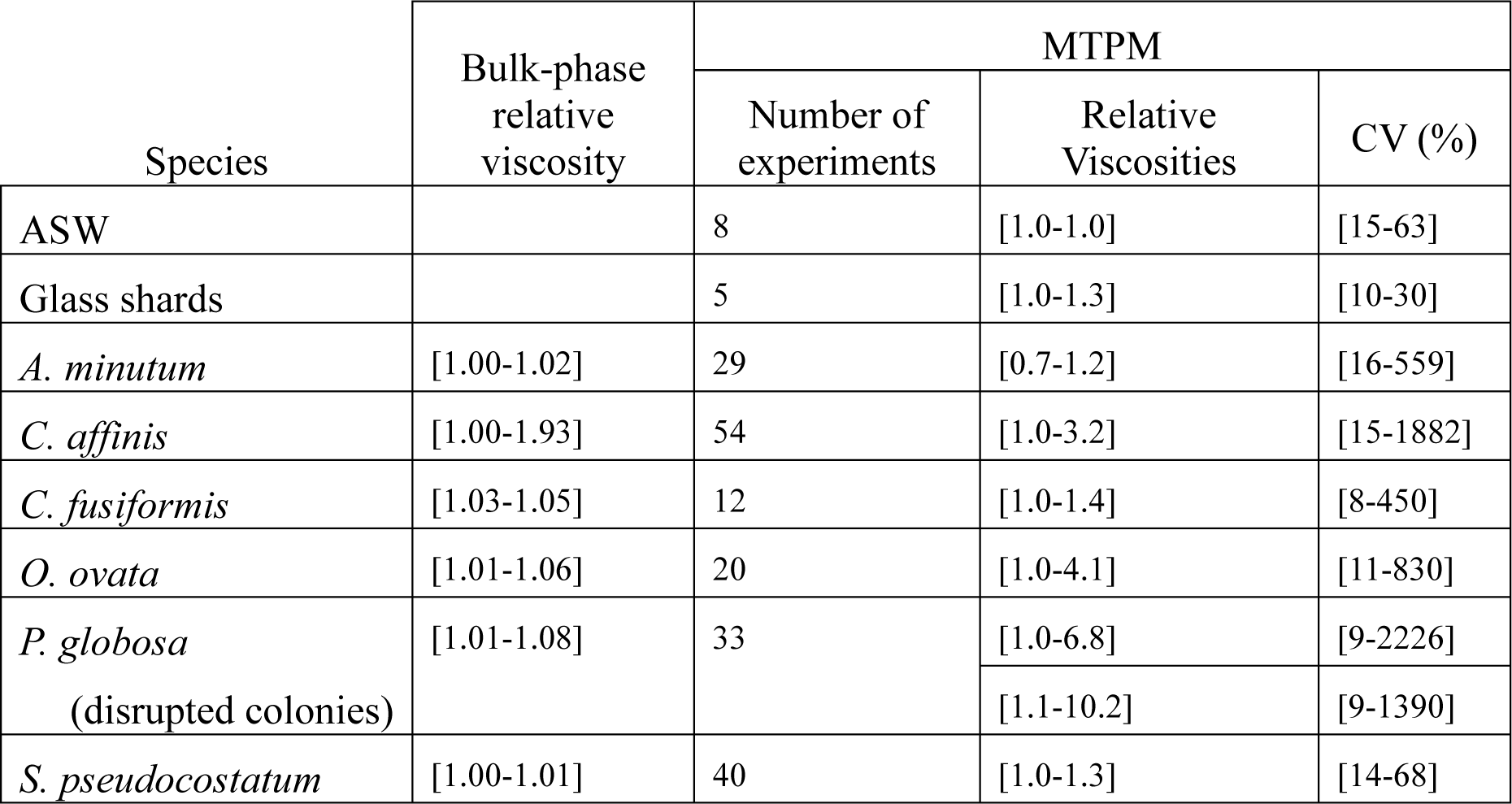
Ranges of bulk-phase relative viscosities, measured with the cone and plate viscometer, and microscale relative viscosities measured with MPTM. CV refers to the coefficient of variability of the 2D viscosity maps.

